# Modeling sensory-motor decisions in natural behavior

**DOI:** 10.1101/412155

**Authors:** Ruohan Zhang, Shun Zhang, Matthew H. Tong, Yuchen Cui, Constantin A. Rothkopf, Dana H. Ballard, Mary M. Hayhoe

**Affiliations:** Department of Computer Science, The University of Texas at Austin, Austin, TX, USA; Computer Science and Engineering, University of Michigan, Ann Arbor, MI, USA; Center for Perceptual Systems, The University of Texas at Austin, Austin, TX, USA; Cognitive Science Center and Institute of Psychology, Technical University Darmstadt, Darmstadt, Germany

## Abstract

Although a standard reinforcement learning model can capture many aspects of reward-seeking behaviors, it may not be practical for modeling human natural behaviors because of the richness of dynamic environments and limitations in cognitive resources. We propose a modular reinforcement learning model that addresses these factors. Based on this model, a modular inverse reinforcement learning algorithm is developed to estimate both the rewards and discount factors from human behavioral data, which allows predictions of human navigation behaviors in virtual reality with high accuracy across different subjects and with different tasks. Complex human navigation trajectories in novel environments can be reproduced by an artificial agent that is based on the modular model. This model provides a strategy for estimating the subjective value of actions and how they influence sensory-motor decisions in natural behavior.

**Author summary:** It is generally agreed that human actions can be formalized within the framework of statistical decision theory, which specifies a cost function for actions choices, and that the intrinsic value of actions is controlled by the brain’s dopaminergic reward machinery. Given behavioral data, the underlying subjective reward value for an action can be estimated through a machine learning technique called inverse reinforcement learning. Hence it is an attractive method for studying human reward-seeking behaviors. Standard reinforcement learning methods were developed for artificial intelligence agents, and incur too much computation to be a viable model for real-time human decision making. We propose an approach called modular reinforcement learning that decomposes a complex task into independent decision modules. This model includes a frequently overlooked variable called the discount factor, which controls the degree of impulsiveness in seeking future reward. We develop an algorithm called modular inverse reinforcement learning that estimates both the reward and the discount factor. We show that modular reinforcement learning may be a useful model for natural navigation behaviors. The estimated rewards and discount factors explain human walking direction decisions in a virtual-reality environment, and can be used to train an artificial agent that can accurately reproduce human navigation trajectories.

## 1 Introduction

Modeling and predicting visually guided behavior in humans is challenging. In various contexts, it is unclear what information is being acquired and how it is being used to control behaviors. Empirical investigation of natural behavior has been limited, largely because it requires immersion in natural environments and monitoring of ongoing behavior. However, recent technical developments have allowed more extensive investigation of visually guided behavior in natural contexts [1]. At the empirical level it appears that complex behaviors can be broken down into a set of subgoals, each of which requires specific visual information [2-4]. In a complex task such as crossing a road, a person must simultaneously determine the direction of heading, avoid tripping over the curb, locate other pedestrians or vehicles, and plan for future trajectory. Each of these particular goals requires some visual evaluation of the state of the world in order to make an appropriate action choice in the moment. A fundamental problem for understanding natural behavior is thus to be able to predict which subgoals are currently being considered, and how these sequences of visuomotor decisions unfold in time.

A theoretical basis for modeling such behavioral sequences is reinforcement learning (RL). Since the breakthrough work by [5], a rapidly increasing number of studies have used a formal reinforcement learning framework to model reward-seeking behaviors. Numerous studies have linked sensory-motor decisions to the underlying dopaminergic reward machinery [1,6]. The basic mechanisms of reinforcement learning, such as reward estimation, temporal-difference error, model-free and model-based learning, and discount factor, have been linked to a broad range of brain regions [7-16]. Because studies of the neural circuitry involve very restrictive behavioral paradigms, it is not known how these effects play out in the context of natural visually guided behavior. Similarly, the application of RL models to human behavior has been restricted almost exclusively to simple laboratory paradigms, and there are few formal attempts to model natural behaviors [17]. The goal of the presented work is to predict action choices in a virtual walking setting by estimating the subjective value of some of the sub-tasks that the sensory-motor system must perform in this context. We show that it is possible to estimate the subjective reward values of behaviors such as obstacle avoidance and path following, and accurately predict the trajectories walkers take through the environment. This demonstration suggests a potential analytical tool for the exploration of natural behavioral sequences.

### Modular reinforcement learning for modeling natural behaviors

The primary focus of reinforcement learning has been on forward models that, given reward signals, can learn to produce policies, which specify action choices when immersed in an environment state. A *state* refers to information about the environment that is needed for decision making. An important breakthrough of RL in behavior modeling is inverse reinforcement learning (IRL), which aims to estimate the underlying subjective reward of decision makers given behavioral data [18]. IRL is an appealing tool for modeling human behavior: A behavioral model can be quantitatively evaluated by comparing human behaviors with reproduced behaviors by an artificial agent trained using the RL model with the estimated reward function.

An important factor that makes standard RL difficult in modeling natural behaviors is its sophistication and resulting computational burden as a model for general reward-seeking behaviors. The natural environment has at least two features that could make RL/IRL algorithms computationally intractable. First, a large number of task-relevant objects may be present, hence the decision state space is likely to be high-dimensional. Standard RL suffers from the *curse of dimensionality* with high-dimensional state space, where the computational burden grows exponentially with the number of state variables [5,19]. Second, the natural environment is ever-changing such that humans must make decisions under different situations although these situations might have similar components. Living in a natural environment requires a decision maker to be able to *transfer* knowledge learned from previous experience to a new situation. In contrast an RL agent is often trained and tested repeatedly in a fixed environment. The optimal behavior is obtained through either a model-based dynamic programming approach that requires full knowledge of the environment, or a model-free learning approach that requires a large amount of experience. Both approaches generally put a heavy burden on memory storage or computation in order to calculate the optimal behavior. Consequently both of them may not be suitable for the real-time decision-making strategy in natural conditions since decision makers encounter new environment all the time and need to make decisions with reasonable cognitive load. For these reasons, standard RL must be extended to make computation tractable.

An extension of standard RL named *modular* reinforcement learning utilizes divide-and-conquer as an approximation strategy [19-21]. The modular RL takes the statistical structure present in the environment, decomposes a task into *modules* where each module solves a subgoal of the original task. Generally an arbitrator is required to synthesize module policies and make final decisions. Modularization alleviates the problem of curse of dimensionality since each module only concerns a subset of state variables. Introducing a new state variable may not affect the entire state space and cause its size to grow exponentially. Additionally, the decomposition naturally allows the decision maker to learn a behavior specifically for a module and reuse it later in a new environment. Under the modular RL framework, a more sample-efficient IRL algorithm is possible [19], which matters for modeling natural human behaviors since such behavioral data is often expensive to collect.

Recent studies have explored the plausibility of a modular architecture for natural visually guided behavior where complex tasks can be broken down into concurrent execution of modules, or microbehaviors [4,9,22,23]. Thus in the example of walking across the street, each particular behavioral subgoal such as avoiding obstacles can be treated as an independent module. This leads to a view of the human brain as the centralized arbitrator that divides and coordinates these modules in a hierarchical manner. The current investigation explores the modular architecture in more detail.

### Estimating the discount factor

A frequently overlooked variable in RL is the discount factor that determines how much a decision-maker weighs future reward compared to immediate reward. In the agent-environment interaction paradigm, a standard RL model typically treats the discount factor as a part of the environment and as fixed. The alternative approach is to view the discount factor as a subjective decision-making variable that is part of the agent and may vary. Behavioral neuroscience studies suggest that the magnitude of the discount factor is correlated with serotonin level in human subjects [24]. As a consequence decision-makers may exhibit between-subject variations [25]. At the same time, between-task variation may also exist, i.e., the same decision maker may use different discount factors for various tasks. An fMRI study by [16] suggests that different cortico-basal ganglia loops are responsible for reward prediction at different time scales, allowing multiple discount factors to be implemented. Hence it is necessary to extend the standard RL model to adapt discount factors to different human subjects and tasks. A modular approach is ideal for this modeling effort. Allowing different modules to have their own discount factors makes the model flexible in modeling potential variations in human data.

Spatial navigation has been used as a canonical benchmark task for standard RL/IRL algorithms in machine learning, and therefore is selected as the experimental domain for testing our model. The task is an ideal testbed for modular RL since it is convenient for introducing multiple (sub-)tasks. In following sections of this paper, computer simulations are conducted first to validate the correctness of the proposed algorithm and to compare with existing methods. We then use human behavioral data previously collected in an immersive virtual environment [4] to show that the proposed sparse modular IRL algorithm allows prediction of human walking trajectories by estimating the subjective reward values and discount factors of different modules. By demonstrating the ability to model naturalistic human sensory-motor behavior we lay the ground work for future analysis of similar behaviors.

## 2 Methods

We introduce the experimental designs and computational models first since they are necessary to understand the results.

### 2.1 Experiments

Virtual reality (VR) and motion tracking were employed to create a naturalistic environment with a rich stimulus array, while maintaining experimental control. Fig 1 shows the basic setup. The subject wore a binocular head-mounted display (the nVisor SX111 by NVIS) that showed a virtual room (8.5 × 7.3 meters). The subject’s eye, head, and body motion were tracked while walking through the virtual room. Subjects were recruited from a subject pool of undergraduates at the University of Texas at Austin, and were naive to the nature of the experiment. The human subject research is approved by the University of Texas at Austin Institutional Review Board with approval number 2006-06-0085 [4].

**Fig 1.**
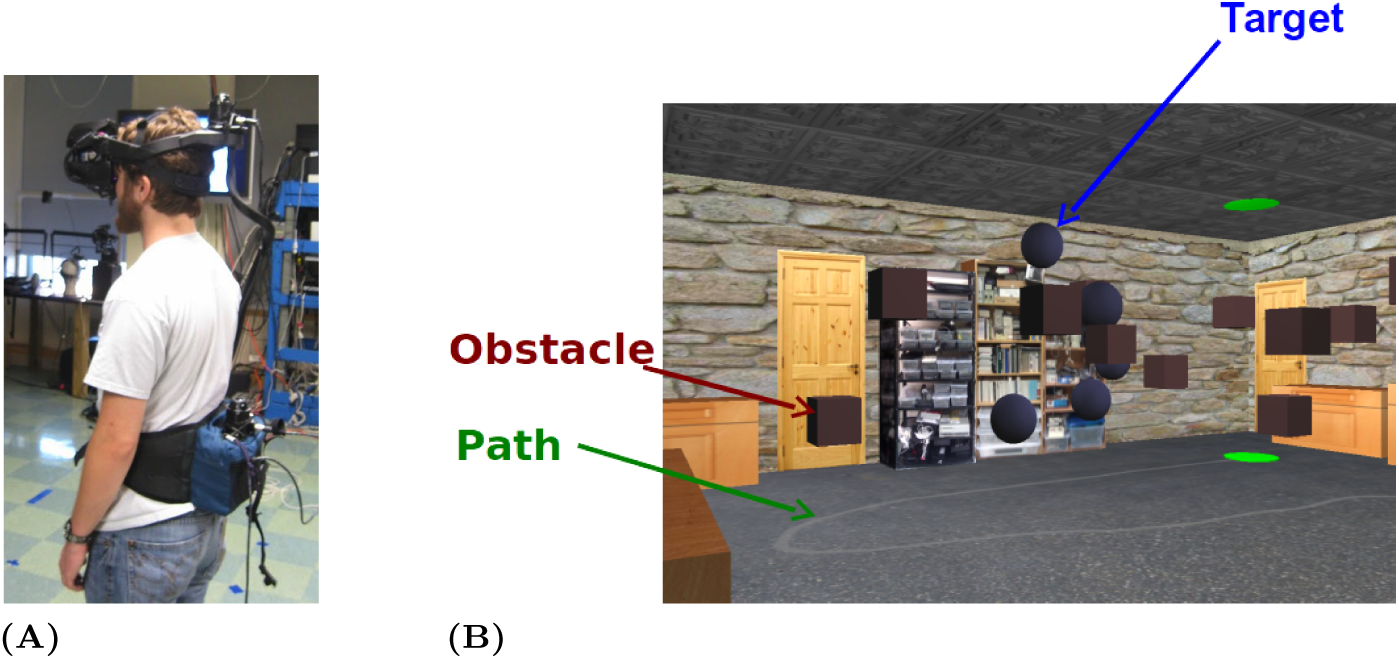
The virtual-reality human navigation experiment with motion tracking. (A) A human subject wears a head mounted display (HMD) and trackers for eyes, head, and body. (B) The virtual environment as seen through the HMD. The red cubes are obstacles and the blue spheres are targets. There is also a gray path on the ground leading to a goal (the green disk). At the green disk the subject is ‘transported’ to a new ‘level’ in a virtual elevator for another trial with a different arrangement of objects.

Although we do not know the set of normal subtasks involved in walking through a room like this, three plausible candidates might be following a path across the room, avoiding obstacles, and perhaps heading towards target objects. To capture some of this natural behavior we asked subjects to collect the targets (blue spheres) by intercepting them, follow the path (the gray line), and/or avoid the obstacles (red cubes). Objects disappeared after collision. This type of state transition function encourages subjects to navigate through the virtual room instead of sticking at a single target.

The global task has at least three *modules*: following the path, collecting targets, and avoiding obstacles. We gave subjects four types of instructions that attempt to manipulate their reward functions (and potentially the discount factors), resulting in four experimental task conditions:

1. **Task 1:** Follow the path only

2. **Task 2:** Follow the path and avoid the obstacles

3. **Task 3:** Follow the path and collect the targets

4. **Task 4:** Follow, avoid, and collect together

There were no monetary rewards in the task. Since following paths, avoiding obstacles, and heading towards targets are frequent natural behaviors, we assume that subjects have some learned, and perhaps context-specific subjective values associated with the three task components, and our goal was to modulate these intrinsic values using the instructions. The instructions were to walk normally, but to give some priority to the particular task components in the different conditions. To encourage such prioritization, Subjects received auditory feedback when colliding with obstacles or targets. When objects were task-relevant, this feedback was positive (a fanfare) or negative (a buzzer), while collisions to task-irrelevant objects resulted in a neutral sound (a soft bubble pop) [4]. The color of the targets and obstacles was counterbalanced in another version of the experiment and was found not to affect task performance or the distribution of eye fixations so the control was not repeated in the present experiment [26]. The order of the task was Task 1, 2, 3, and 4. This order was chosen so as not to influence the single task conditions by doing the double task. Thus it is possible there are some order effects. In another experiment in the environment the order of the conditions was counterbalanced and no obvious order effects were observed [26].

We analyze data collected from 25 human subjects. A single experimental trial consisted of a subject traversing the room, with the trial ending when the goal at the end of the path is reached. Objects’ positions and the path’s shape differed on every trial. Each subject performed four trials for each task condition.

#### Data availability

This general paradigm of navigation with targets and obstacles has been used to evaluate modular RL and IRL algorithms [2,19] and to study human navigation and gaze behaviors [4,27]. The data that support the findings of this study are made public and available at [28].

### 2.2 Modular Reinforcement Learning

Reinforcement learning basics A standard reinforcement learning model is formalized as a Markov decision process (MDP). The MDP models the interaction between the environment and a decision maker which will be referred as an agent. Formally, an MDP is defined as a tuple ⟨*𝒮, 𝒜, 𝒫, ℛ, γ*⟩ [5], where:

- 𝒮 is a finite set of environment states. Let *s*_*t*_ denote the agent’s state at discrete time step *t*. The state encodes relevant information for an agent’s decision.
- 𝒜 is a finite set of available actions. Let *a*_*t*_ be the action agent chooses to take at time *t*. The agent interacts with the environment by taking an action in its observed state.
- 𝒫 is the state transition function which specifies the probability P(*s*’|*s*, a), i.e., the probability of entering state *s*’ when agent takes action *a* in state s. The state transition function describes the dynamics of the environment that are influenced by an agent’s action.
- ℛ is a reward function. *r_t_* denotes the scalar reward agent received at time step *t*.
- γ ε [0,1) is a discount factor. The agent values future rewards less than an immediate reward, therefore future rewards are discounted by parameter γ at every discrete time step. γ = 0 indicates that the agent is myopic and only seeks to maximize the immediate reward.
- π : 𝒮 ↦ 𝒜 is called a policy of the agent, which specifies the probability of chosen each action in each state.

In machine learning, the purpose of a reinforcement learning algorithm is to find an optimal policy π* that maximizes the longterm cumulative reward. Many RL algorithms are based on value function estimation. The action-value function (also called Q-value function) estimates the expected longterm reward for taking an action in a given state, and follow policy π afterwards. Formally, the Q-value function conditioned on policy π is defined as [5]:

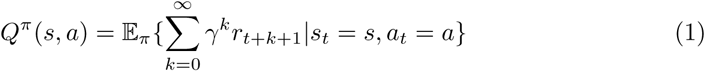

Given the Q-value function it is convenient for an agent to select the action that maximizes expected future returns.

#### Modular Reinforcement Learning

The divide-and-conquer approximation of RL leads to modular reinforcement learning, in which a *module* is a subtask of the original task. Each module is hence a simpler problem, so that its value function and policy can be learned or calculated efficiently. A module is also modeled by an MDP 𝒮 ^(*n*)^, 𝒜, 𝒫 ^(*n*)^, ℛ ^(*n*)^, γ ^(*n*)^, where n is the index of the nth module. Note that each module has its own state space, transition function, reward function, and discount factor, but the action space is shared between modules because all modules reside in a single agent.

Let *N* be the number of modules and 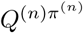 denote module Q-value function of the *n* th module conditioned on module policy π ^(*n*)^. For simplicity, we will drop π ^(*n*)^ and write Q^(*n*)^. Let Q without superscription denote the global Q function (also drop global policy π). Modular RL sums module Q functions to obtain the global Q function [21,29]:

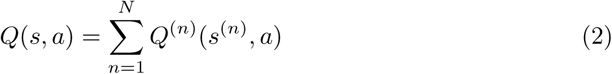

There can be multiple *module objects* of a module, e.g., several identical obstacles nearby to avoid. The number of objects of each module is denoted as *M*^(1)^,…, *M*^(*N*)^. Note that for a given module, its module objects share the same Q^(*n*)^ since their module MDPs are identical. But at a given time they could be in different states relative to the agent’s reference frame which can be denoted as *s^(n,m)^* for module n object m. To generalize the above equation:

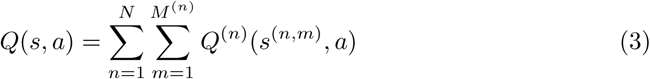

This assumes independent transition functions between module objects [19]. A module action-value function Q^(*n*)^ may be calculated from solving Bellman equations using dynamic programming or through standard learning algorithms with enough experience data, which we argue to be infeasible for human performing natural tasks. Q^(*n*)^ needs to be calculated efficiently with reasonable cognitive load.

In our experiments, both the state transition function and reward function are deterministic hence the expectation in Eq (1) can be dropped. Since each module Q function only considers a single source of reward from a single module object, and assuming a policy that leads the agent directly to the module object, Q^(*n*)^(*s*^(*n,m*)^, *a)* takes the following simple form:

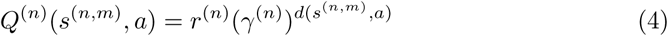

where r^(*n*)^ is the reward for the nth module, γ ^(*n*)^ is its discount factor, and *d*(*s*^(*n,m*)^, *a)* is the spatial or temporal distance between the agent and the module object *m* after taking action *a* at state *s*^(n,m)^. Note Eq (4) converts value function back to its simplest form in [15]. This simple form allows a decision maker to calculate the action-value for a state efficiently when needed instead of beforehand. This matters when humans need to make decisions fast and when it is computationally expensive to calculate value functions using a standard RL algorithm. It is also unlikely for a human to pre-compute the values for all future states and use dynamic programming to obtain a global policy when they visit the environment for the first time. Doing so would at least require a human to store Q-values for relevant states (a Q-table) in its memory system, which is convenient for an artificial agent but would be difficult for a real-time human decision maker.

Why does modular RL alleviate the problem of curse of dimensionality? Consider the joint state space of a standard RL which can be represented as the Cartesian product of the module state spaces: 𝒮 = 𝒮 ^(1)^ x 𝒮 (^2^) x …. The computation cost for one iteration in value iteration (a popular RL algorithm) is *O* (| 𝒮 |^2^ | 𝒜|) where |. | denotes the cardinality of a set [30]. When a new module S^(N^) is added, the cost of standard RL becomes *O* (|S^(1)^ x 𝒮 ^(2)^ x … x 𝒮 ^(N^)|^2^|𝒜|), while the cost of modular RL becomes *O* (| 𝒮 ^(1)^|^2^|𝒜|) + *O* (| 𝒮 ^(2)^|^2^| 𝒜|) + … + *O* (|S^(*n*)^|^2^| 𝒜|). Therefore the computational cost increases additively in modular RL instead of multiplicatively.

#### Visualizing modular reinforcement learning

Eq (4) bridges modular RL with an important planning method called artificial potential field [31-33]. Similar to a potential field, we use a value surface to visualize the value function. Each module objects is associated with a value surface. The module reward controls the maximum absolute height of the surface, and the discount factor controls temporal or spatial discounting rates. Module value surfaces can be composed directly by summation or integration to produce a multi-module value surface. The concept of value surfaces and their combination is illustrated in Fig 2. Given a composed value surface as in Fig 2F, a modular RL agent would choose actions that lead to a local minima on the surface. A sequence of actions could construct a trajectory in Fig 3A which traverses through a sequence of local minima.

**Fig 2.**
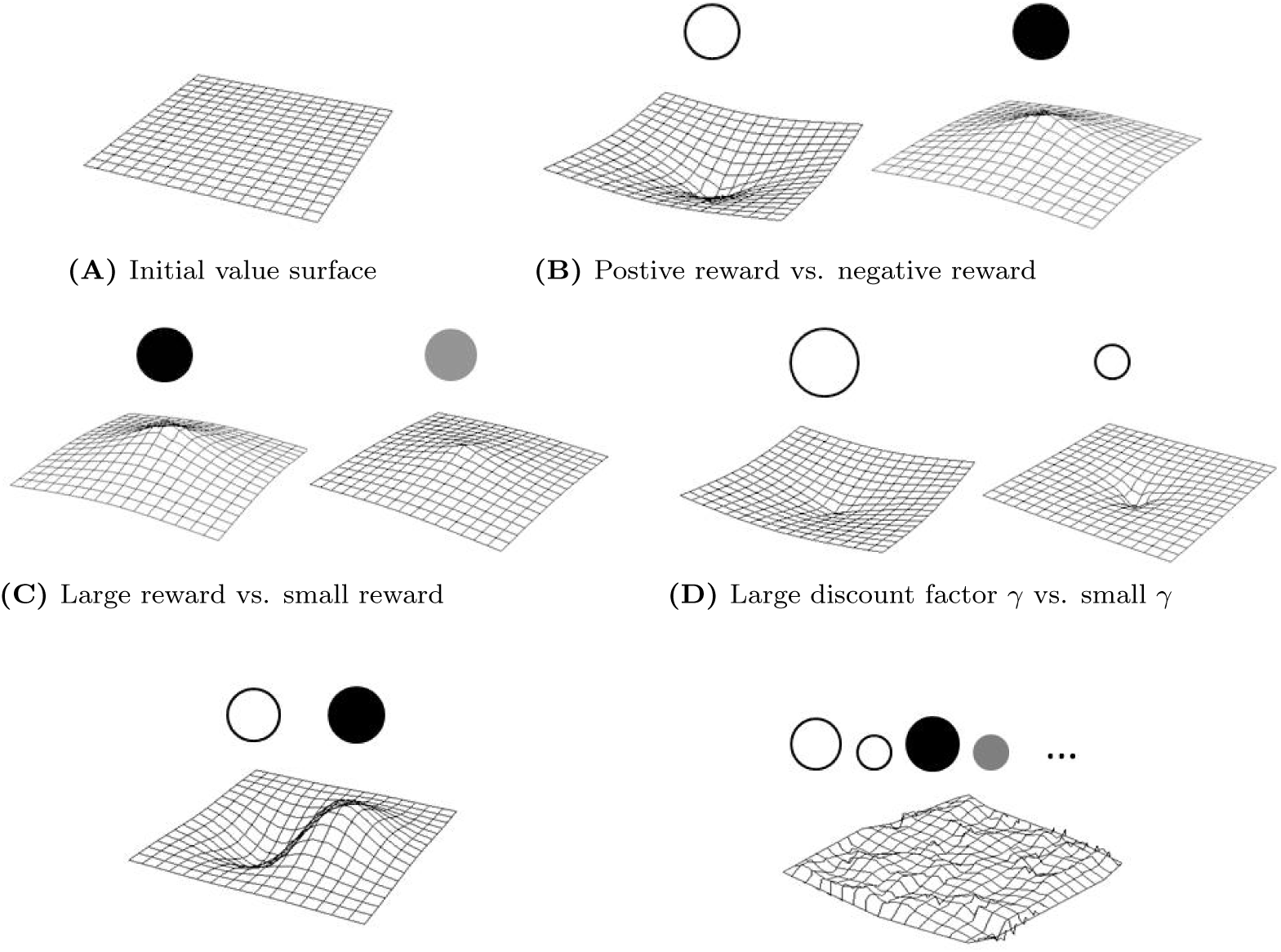
The concept of modular reinforcement learning illustrated using value surfaces. (A) The value surface is flat without any reward signal. (B) A module object with positive reward has positive weight, and one with negative reward has negative weight. They bend the value surface to have negative and positive curvatures respectively. Therefore, an agent desires to follow the steepest descent to minimize energy, or equivalently, to maximize reward. (C) An object with larger weight bends the surface more. (D) An object with greater discount factor γ has larger influence over distance. (E,F) Composing different objects with different rewards and γ_S_ results complicated value surfaces that can model an agent’s value function over the entire state space.

**Fig 3.**
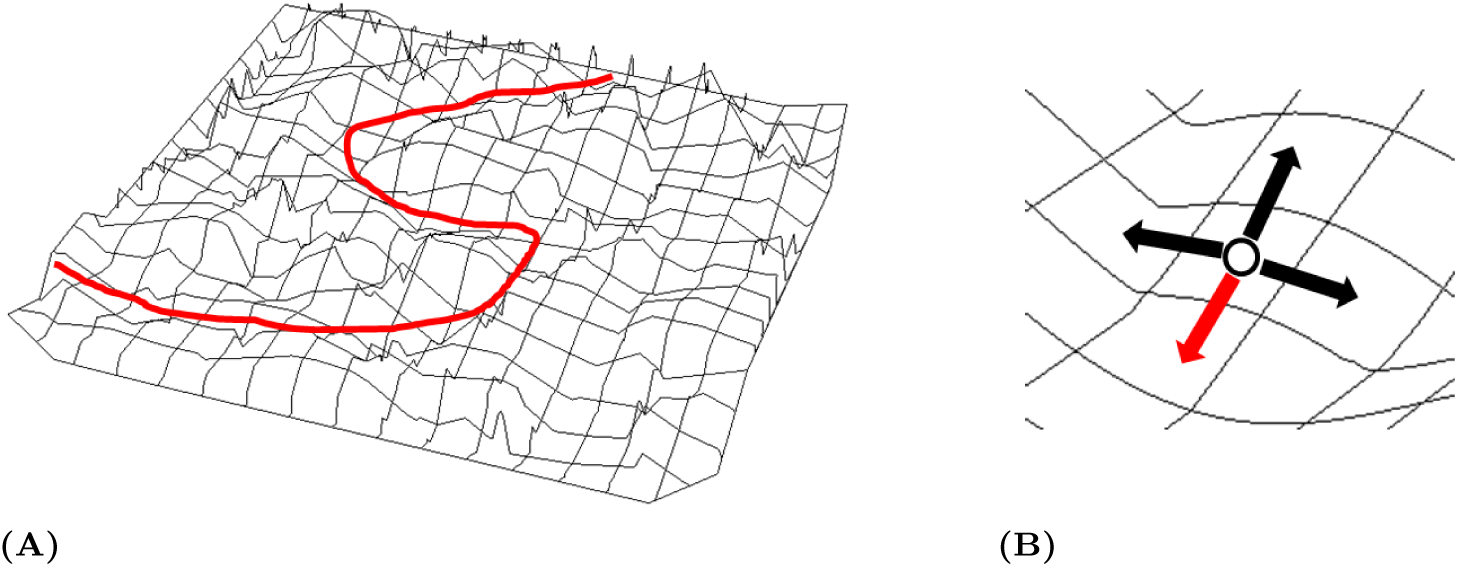
Maximum likelihood modular inverse reinforcement learning. (A) From an observed trajectory (a sequence of state-action pairs), the goal of modular IRL is to recover the underlying value surface. (B) Maximum likelihood IRL assumes that the probability of observing a particular action (red) in a state is proportional to its Q-value among all possible actions as in Eq (5).

### 2.3 Modular Inverse Reinforcement Learning

While reinforcement learning aims at finding the optimal policy given a reward function, inverse reinforcement learning (IRL) attempts to infer the unknown reward function given the agent behavioral data in the form of state-action pairs (*s*_*t*_, *a*_*t*_) [18,34-36]. Our work is largely based on the modular IRL algorithm by [19] which pioneered the first modular IRL algorithm. Given the modular RL formulation in the previous section, the goal of modular IRL is to estimate the underlying reward and discount factor for each module to recover the value function, given a sequence of observed state-action pairs, i.e., a trajectory that traverses through the state space, as shown in Fig 3A.

We follow the Bayesian formulation of IRL [36,37], Maximum Likelihood IRL [38], and improve the modular IRL algorithm in [19]. These approaches assume that the higher the Q-value for an action *a*_*t*_ in state s_t_, the more likely action *a*_*t*_ is observed in behavioral data. Let *η* denote the confidence level in optimality (the extent to which an agent selects actions greedily, default to be 1), and let exp(·) denote the exponential function. The likelihood of observing a certain state-action pair is modeled by the softmax function with Gibbs (Boltzmann) distribution, as illustrated in Fig 3B:

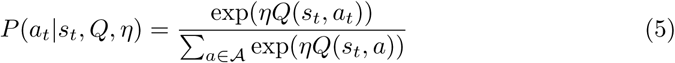

Let *T* denote the total length of the trajectory. The overall likelihood ℒ for observed data D = {(*s*_1_, *a*_1_), …, (*s*_*T*_, *a*_*T*_)} is the product of the likelihood of individual state-action pairs, given the states are Markovian and action decisions are independent:

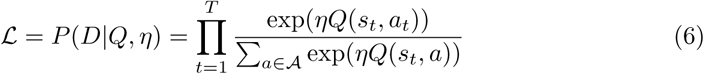

Next, the global action-value function *Q*(*s*_*t*_, *a*_*t*_) is decomposed using Eq (3) with module Q functions *Q*^(*1:N*)^, therefore the likelihood becomes:

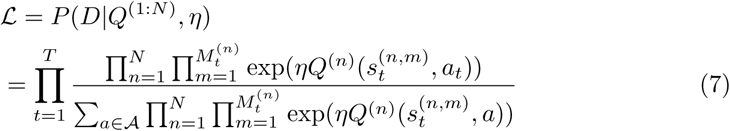

Take the log of the likelihood function:

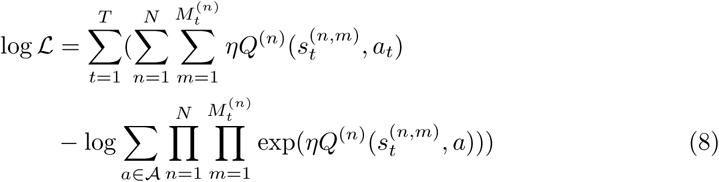

Substituting Eq (4) into Eq (8):

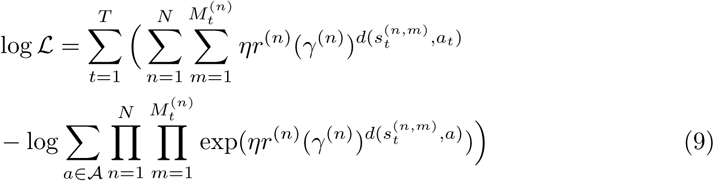

The variables to be estimated from the data are module rewards *r*^(*1:N*)^ and discount factors γ ^(*1:N*)^ The number of modules N, the number of objects for each module 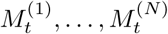 and distances 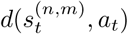 for each object are all state information and can be observed from the environment. This formulation follows closely the work by [19], extending it to use the new formulation of modular RL, handle multiple objects of each module, estimate the discount factors, and derive a slightly different objective function.

#### Sparse modular inverse reinforcement learning

Modular IRL can only guess which objects are actually being considered by the decision maker when chosen an action. To address this problem, we can further add a *L*_*1*_ regularizer 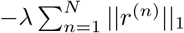 to Eq (9), which causes some module rewards to become 0 so these modules would be ignored in decision making. This is an extension of using a Laplacian prior in Bayesian IRL [36]. In addition to the benefit from an optimization perspective, the regularization term has the following important interpretation in terms of explaining natural behaviors.

A *hypothetical module set* is a set ℋ = {1, …, *N*} contains *N* modules that could potentially be of an agent’s interest. However, due to the limitations in computational resource, the agent can only consider a subset of ℋ at a time, denoted ℋ’. In a rich environment many modules’ rewards would be effectively zero at current decision step, hence | ℋ’| ≪| ℋ|. For instance, a driving environment could contain hundreds of objects in ℋ. But a driver may pay attention to only a few. The regularization constant λ serves as a cognitive capacity factor that helps determine ℋ from the observed behaviors. Therefore the final objective function of modular IRL is:

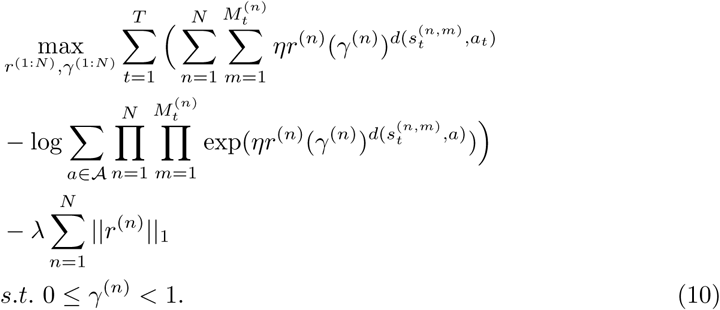

Note that if we are to fit *r*^(*1:N*)^ and *γ* ^(*1:N*)^ simultaneously, the above objective function is non-convex. However, the objective becomes convex if only fitting *r*^(*1:N*)^. Since *γ* ^(*n*)^ is in range (0,1), one can perform a grid search over values for *γ* ^(*1:N*)^ with step size e and fit *r*^(*1:N*)^ at each possible *γ* ^(*1:N*)^ value. This allows us to find a solution within ε-precision of the true global optimum.

An accessible evaluation of the proposed algorithms in an artificial multitask navigation environment can be found in Appendix 1. The environment is a 2D gridworld that resembles the virtual room we use for the human experiments. The validity of the modular IRL is proved empirically by showing its ability to recover true module rewards and discount factors with high accuracy given enough behavioral data. Meanwhile it requires significantly less data samples to obtain high prediction accuracy comparing to a standard Bayesian IRL algorithm [36], presumably because the state space is reduced significantly by modularization. Sparse modular IRL is shown to further improve sample efficiency if task-irrelevant modules are present. Unlike computer simulated experiments in which one can easily generate millions of behavioral data, human experiments have a more expensive data collection procedure in general. Therefore sample efficiency of sparse modular IRL is an important advantage in modeling natural human behaviors, which will be seen in the next section.

## 3 Results

Despite its computational advantages shown in simulation, the question remains whether modular IRL can be used as a decision-making model to explain human behaviors in the experiments. Sparse modular IRL (Eq (10)) is used as the objective function to estimate reward *r* and discount factor *γ* for the target, obstacle, and path modules. However the regularization constant is found to be close to zero since there are only three modules. Recall that each subject performs each task four times, and each time the path and the arrangement of objects are different. We use leave-one-out cross evaluation, where *r, γ* are estimated using all-but-one training trials that are from the same subject and same task condition and evaluated on the remaining test trial. Since the parameter estimates are based on the other three trials, all of our prediction results shown below are for a *novel* environment with similar components - this requires the model to generalize across environments. The number of data samples obtained from a single trial is typically around 100 hence sample efficiency is critical for the performance of an algorithm.

Different *r* and *γ* are estimated for each subject under each task condition for each module, hence there are 25 subjects × 4 conditions × 3 modules × 4 trials = 1,200 different pairs of *r, γ* estimations. The state information for the model includes the distance and angle to the objects, while the state space is discretized using grids of size 0.572 by 0.572 meters, a parameter chosen empirically that produces the best modeling result. It also matches the approximate length of a step in VR, so is a suitable scale for human direction decisions. Empirically, as long as the grid size is within reasonable range of human stride length (0.3-0.9 meters) the algorithm’s performance is fairly robust.

The path is discretized into a sequence of waypoints which are removed after being visited (similar to the targets). The action space spans 360 degrees and is discretized to be 16 actions using bins of 22.5 degrees. This is a suitable discretization of the action space, given the size of the objects at the distance of 1-2 meters, where an action decision is most likely made.

### Qualitative results and visualization

The most intuitive way to evaluate the modular RL model is to see whether the model can accurately reproduce human navigation trajectories. The Q-value function of a modular RL agent is calculated using *r* and *γ* estimated from human data. Next, the modular RL agent is placed at the same starting position as the human subject and starts to navigate the environment until it reaches the end of the path. The agent chooses an action probabilistically based on the Q-value of the current state, using a softmax action selection function as in Eq (5). The reason to let the agent choose actions with a certain degree of randomness is that the Q-values for multiple actions can be very close, e.g., turning left or turning right to avoid an obstacle, consequently a human subject may choose either. Therefore, a single greedy trajectory may not overlap with the actual human trajectory. The softmax action selection function generates a distribution of hypothetical trajectories, i.e., a trajectory cloud, by running an agent many times in the same environment. The actual human trajectory can be visualized in the context of this distribution.

Fig 4 shows generated trajectory clouds together with actual human trajectories, along with estimated rewards and discount factors. The agent trajectories are shown in semi-transparent green hence darker area represents trajectories with higher likelihood, and the human trajectory on that trial is shown in black. Each row of figures presents experimental trials from one experimental condition (Task 1-4), and three trials within each row are from different subjects but the same environment, i.e., the same arrangement of objects.

**Fig 4.**
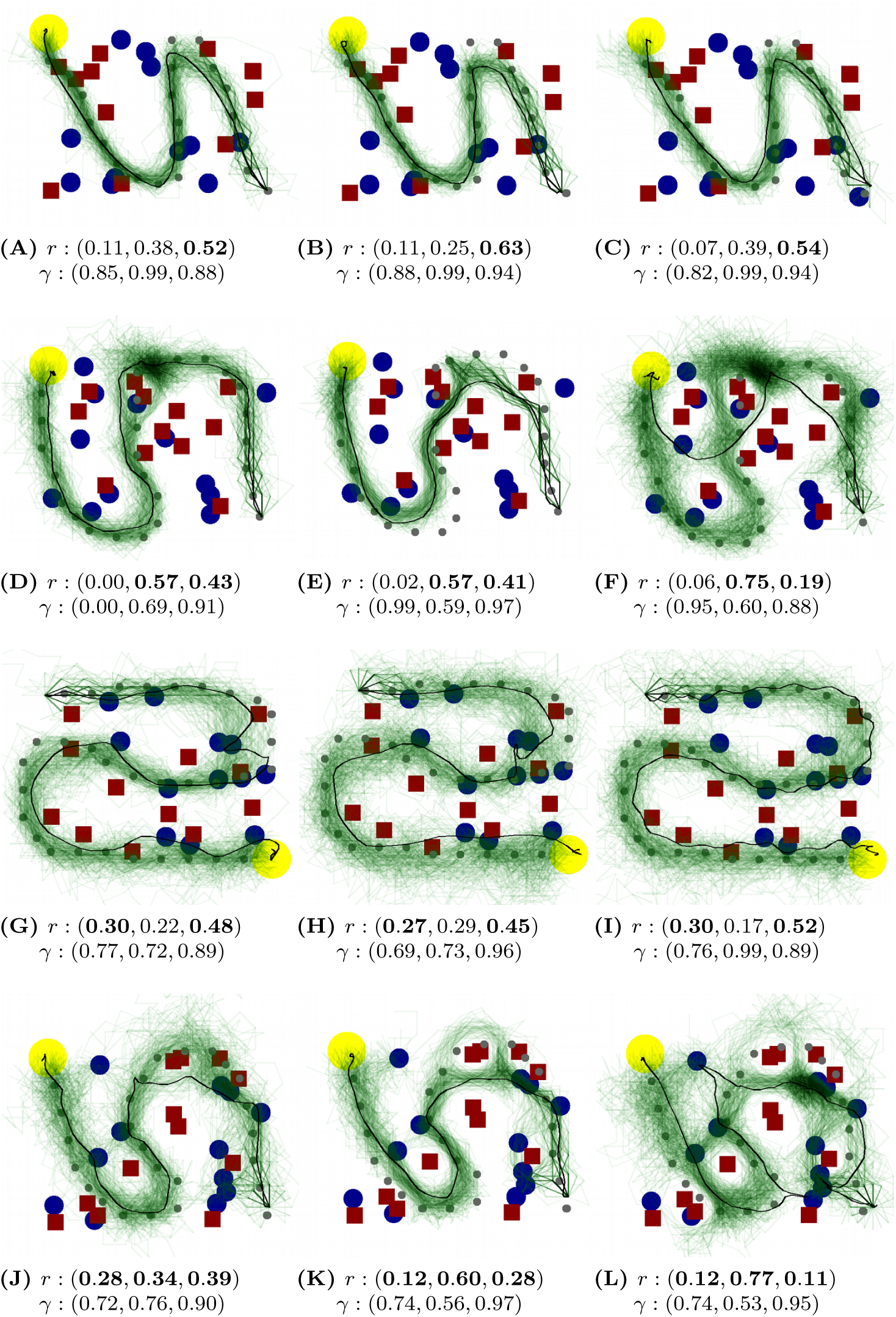
Bird’s-eye view of human trajectories and agent trajectory clouds across different subjects. Black lines: human trajectories. Green lines: modular RL agent trajectory clouds generated using softmax action selection. The green is semi-transparent hence darker area represents trajectories with higher likelihood. Yellow circles: end of the path. Blue circles: targets. Red squares: obstacles. Gray dots: path waypoints used by the model (subjects see a continuous path). Below each graph are the rewards and discount factors estimated from human and used by the modular RL agent. The rewards and discount factors are shown in the order of (Target, Obstacle, Path). The module rewards that correspond to task instructions are bold. Obstacle module has negative reward, but to compare with the other two modules the absolute value is taken. Three trials within each row are from different subjects but the same environment. (A,B,C) show trials from **Task 1: follow the path.** (D,E,F) show trials from **Task 2: follow the path and avoid obstacles**: (G,H,I) show trials from **Task 3: follow the path and collect targets**. (J,K,L) show trials from **Task 4: follow the path, collect targets, and avoid obstacles**.

The figures demonstrate that the model’s generated trajectory clouds align well with observed human trajectories. When a local trajectory distribution is multi-modal, e.g., in Fig 4D, 4F, 4J, 4K, and 4L, the human trajectories align with one of the means. The next important observation is the between-subject variation. Trials within each row are from the same environment under the same task instruction. However, human trajectories can sometimes exhibit drastically different choices, e.g., Fig 4E versus 4F, 4J versus 497.K. These differences are modeled by the underlying *r* and *γ*, and accurately reproduced by the distributions generated. This means that we can compactly model naturalistic, diverse human navigation behaviors using only a reward and a discount factor per module. The modeling power of modular RL is demonstrated by the observation that varying these two variables can produce a rich class of human-like navigation trajectories.

### Between-task and between-subject differences

We then look at the way average reward estimates vary between different tasks when aggregating data from all subjects. The results are shown in Fig 5A. Overall, the estimated *r* values vary in an appropriate manner with task instructions. Thus obstacles are valued higher when the instructions prioritize this task, and targets are valued higher when that task is prioritized. Note that the obstacle avoidance module is given some weight even when it is not explicitly prioritized - this is consistent with the observation that subjects deviates from the path to avoid obstacles even when obstacles are task-irrelevant. This may reflect a bias which is carried over from natural behavior with real obstacles. The relatively high value for the path may indicate that subjects see staying near the path as the primary goal.

**Fig 5.**
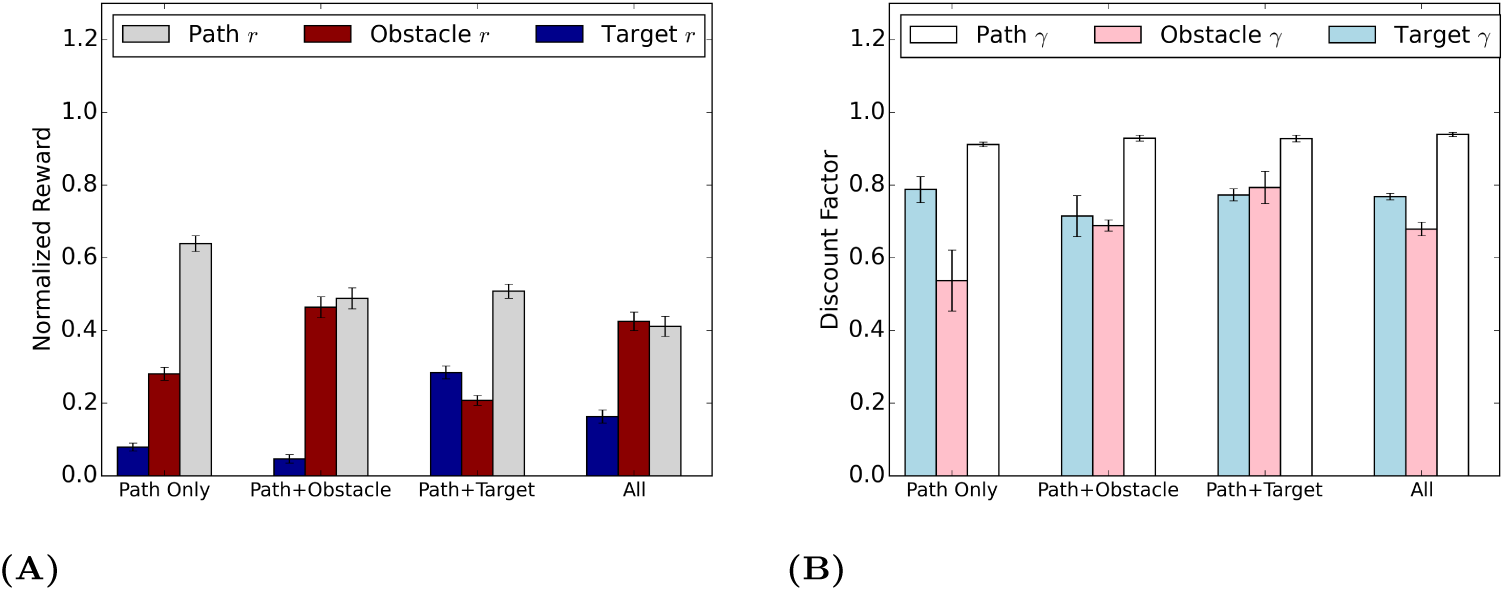
(A) Normalized average rewards across different task instructions. The error bar represents the standard error of the mean between subjects (*N* = 25). The obstacle module has negative reward, but to compare with the other two modules its absolute value is taken. The estimated reward agree with task instructions. (B) Average discount factors across different task instructions. The error bar represents the standard error of the mean between subjects (*N* = 25).

The between-subject differences in reward are shown in Appendix 2 for all 25 subjects. At each individual subject’s level, changing in the relative reward between the modules is also consistent with task instructions. An one-way ANOVA test suggests that individual differences are evident across subjects under the same task instruction (see Appendix 2 for details).

Fig 5B shows average discount factor estimates for different tasks. Although the reward evidently reflects and agrees with task instructions, the interpretation of the discount factor is more complicated. The discount factors vary across tasks for target and obstacle modules but are close to 1.0 and stable for the path module. This may also reflect the primacy of the task of getting across the room, and the need to plan ahead. Although the instructions do not directly manipulate discount factors, we will later show that estimating discount factors from data instead of holding them fixed is important for modeling accuracy.

### Stability of rewards and discount factors across tasks

An important observation from Fig 5 is that *task-relevant* module rewards and discount factors are stable across task conditions. To show this quantitatively, for each subject, we combine module rewards from Task 2 (path + obstacle) and Task 3 (path + target) to synthesize the rewards for Task 4 (path + obstacle + target) in the following way:

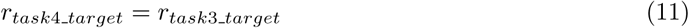

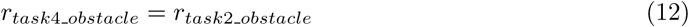

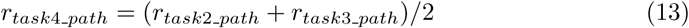

Then the discount factors are synthesized in the similar way. The synthesized rewards (re-normalized) and discount factors from Task 2 and 3 are found to be very close to those estimated from Task 4, as shown in Table 1. However, task-irrelevant rewards and discount factors are not stable. This result indicates that task-relevant module rewards and discount factors generalize to a different task condition. Thus modules are independent and transferable in this particular scenario.

**Table 1.**
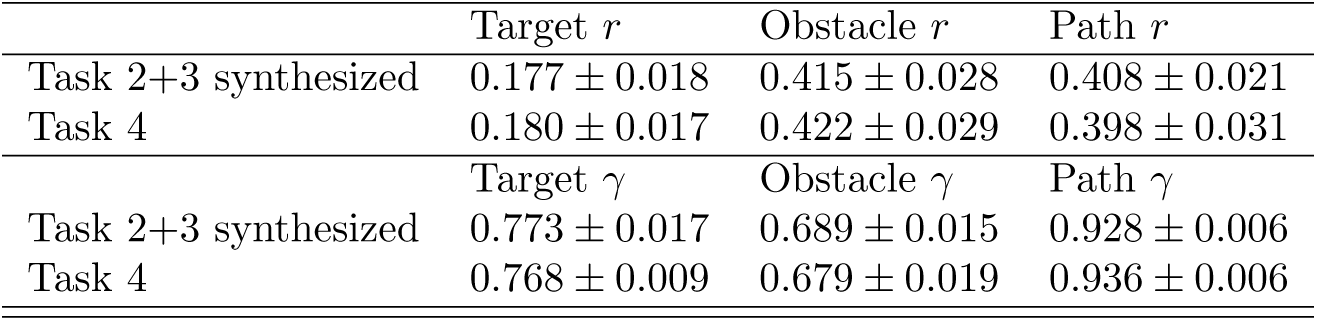
Synthesized rewards and discount factors compared to the estimated ones. Rewards are re-normalized. Results are presented as mean ± standard error between subjects (*N*=25).

| | Target $r$ | Obstacle $r$ | Path $r$ |
| --- | --- | --- | --- |
| Task 2+3 synthesized | $0.177 \pm 0.018$ | $0.415 \pm 0.028$ | $0.408 \pm 0.021$ |
| Task 4 | $0.180 \pm 0.017$ | $0.422 \pm 0.029$ | $0.398 \pm 0.031$ |
| | Target $\gamma$ | Obstacle $\gamma$ | Path $\gamma$ |
| Task 2+3 synthesized | $0.773 \pm 0.017$ | $0.689 \pm 0.015$ | $0.928 \pm 0.006$ |
| Task 4 | $0.768 \pm 0.009$ | $0.679 \pm 0.019$ | $0.936 \pm 0.006$ |

### Quantitative results and comparisons to alternative models

Next we compare our model with several alternative hypotheses. The full modular IRL model chooses the action greedily that maximizes the Q-value function of each state using both estimated *r* and *γ*. An ablation study is conducted to demonstrate the relative importance of the variables in the model. The binary reward agent estimates *γ* only, and uses a unit reward of 1 for the module that is task-relevant, e.g., in Task 2 the path and the obstacle modules would have rewards of +1 and −1 respectively, and the target module would have a reward of 0. The fixed *γ* agents estimate *r* only, and use fixed *γ* = 0.1, 0.5, 0.99. A Bayesian IRL agent without modularization and assumes a fixed discount factor [36] is also implemented where the implementation details can be found in Appendix 3.

We choose two performance metrics to evaluate these models. The first one is the number of objects intercepted by the agent’s entire trajectory under different task conditions. Fig 6 shows the performance of different models ((A) targets and (B) obstacles). Overall, the modular IRL model has the closest performance to the human data across task conditions. Note that the number of targets collected is only a little affected by the avoid instruction and obstacles avoided do not change very much with the target instruction, supporting the previous claim that the modules in this experiment are independent hence task-relevant module values are stable. Bayesian IRL and fixed *γ* = 0.99 models perform poorly-the number of objects hit does not vary accordingly with task instructions. The binary reward model, *γ* = 0.1, 0.5 reflect task instructions correctly but are less accurate than the full modular IRL model.

**Fig 6.**
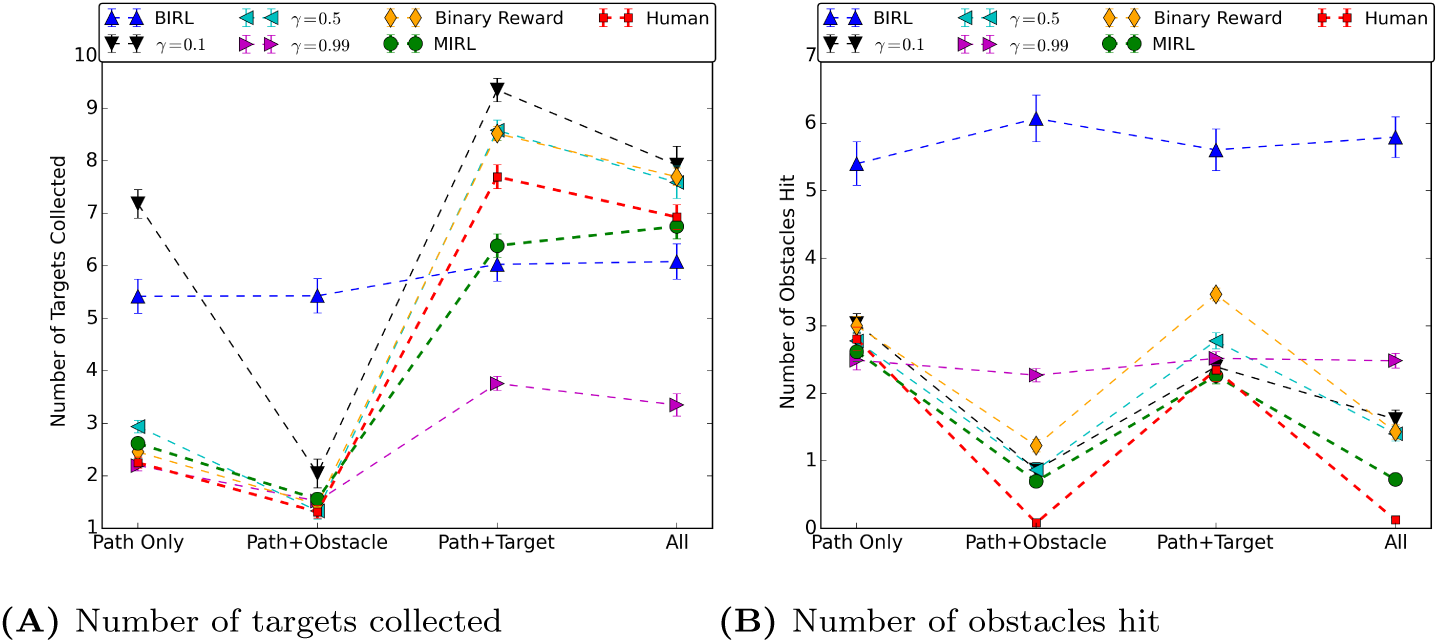
Average number of targets collected/obstacles hit when different models perform the navigation task across all trials. There are 12 targets/obstacles each in the virtual room. Error bars indicate standard error of the mean (N = 100).

The second quantitative evaluation metric would be the angular difference, i.e., policy agreement, which is obtained by placing an agent in the same state as a human and measuring the angular difference between the agent’s action and the human subject’s action. This metric differs from the previous one because it emphasizes more on the accuracy of local decisions instead of the whole trajectory. Thus this angular difference is a local metric instead of a holistic one. The comparison results are shown in Table 2. All modular RL agents are more accurate in predicting human actions comparing to the traditional Bayesian IRL algorithm. Again the full modular IRL model results in higher accuracy comparing to the alternative models. The binary reward model has comparable performance and is in general better than the models that have the discount factor fixed. This supports our claim that module-specific discount factor plays an important role in modeling human behaviors and should be estimated from data.

**Table 2.** Evaluation of the modular agent’s performance compared with baseline agents, measured by the average angular difference (in degrees) compared to actual human decisions. The results are presented as mean ± standard error (N = 100). The agent that uses the full model outperforms all other models.

|  | Task 1 | Task 2 | Task 3 | Task 4 |
| --- | --- | --- | --- | --- |
| Bayesian IRL | 53.87±2.54 | 53.37 ± 2.71 | 59.86 ± 2.00 | 51.09 ± 2.60 |
| Fixed $\gamma = 0.1$ | 31.74±0.88 | 39.43±1.18 | 36.16±0.75 | 41.40±0.88 |
| Fixed $\gamma = 0.5$ | 21.46±0.46 | 36.04±1.16 | 34.20±0.78 | 39.14±0.92 |
| Fixed $\gamma = 0.99$ | 18.19±0.32 | 27.63±1.41 | 28.61±0.93 | 31.63±1.08 |
| Binary Reward | 17.66±0.38 | 27.66±1.44 | 29.97±0.72 | 29.80±0.95 |
| MIRL (Full Model) | 17.94±0.33 | 27.39±1.46 | 26.98±0.80 | 27.65±1.02 |

To summarize, we are able to predict human novel trajectories in different environments on the basis of rewards and discount factors estimated from behavioral data. Since we do not know the actual set of visual operations involved in walking through a cluttered room like this, the fact that we can reproduce the trajectories suggests that the three chosen modules can account for a substantial fraction of the behavior while vision may be used for other tasks. In fact, close to half the fixations made by the subject are on regions of the environment other than the path or objects [4]. This suggests that there may be other visual computations going on but that they do not have much influence on the behavior. Thus the modular RL agents generate reasonable hypotheses about underlying human decision-making mechanism.

These results provides a strong support for using modular RL as the model for explaining such multitask navigation behaviors, and modular IRL as a sample efficient algorithm to estimate rewards and discount factors. Bayesian IRL has to deal with a complex high-dimensional state space and settle for its approximations for a dynamic multi-task problem with limited data, while modular RL can easily reduce the dimensionality of the state-space by factoring out sub-tasks. Therefore the algorithm significantly outperforms the previous standard IRL method in terms of the accuracy in reproducing human behaviors.

## 4 Related Work in Reinforcement Learning

The proposed modular IRL algorithm is an extension and refinement of [19] which introduced the first modular IRL and demonstrated its effectiveness using an simulated avatar. The navigation tasks are similar but we use data from actual human subjects. While they use a simulated human avatar and moving from the straight path, our curved path proves quite different in practice, as well, being significantly more challenging for both humans and virtual agents. We then generalize the state space to let the agent consider multiple objects for each module, while the original work assumes the agent considers one nearest object of each module.

Bayesian IRL was first introduced by [36] as a principled way of approaching an ill-posed reward learning problem. Existing works using Bayesian IRL usually experiment in discretized gridworlds with no more than 1000 states with an exception being the work of [39] which was able to test on a goal-oriented MDP with 20,518 states using hierarchical Bayesian IRL.

The modular RL architecture proposed in this work is most similar to a recent work in [40], in which they decompose the reward function in the same way as the modular reinforcement learning. Their focus is not on modeling human behavior, but rather on using deep reinforcement learning to learn a separate value function for each subtask and combining them to obtain a good policy. Other examples of divide-and-conquer approach in RL include factored MDP [41] and co-articulation [42].

Hierarchical RL [43,44] utilizes the idea of *temporal abstraction* to allow more efficient computation of the policy. [45] analyzes human decision data in spatial navigation tasks and the Tower of Hanoi; they suggest that human subjects learn to decompose tasks and construct action hierarchy in an optimal way. In contrast with that approach, modular RL assumes *parallel decomposition* of the task. The difference can be visualized in Fig *γ*. These two approaches are complementary, and are both important for understanding and reproducing natural behaviors. For example, a hierarchical RL agent could have multiple concurrent *options* [43, 44] executing at a given time for different behavioral objectives. Another possibility is to extend the modular RL to a two-level hierarchical system. Learned module policies are stored and a higher-level scheduler or arbitrator decides which modules to activate or deactivate given the current context and the protocol to synthesize module policies. An example of this type of architecture can be found in [2].

**Fig 7.**
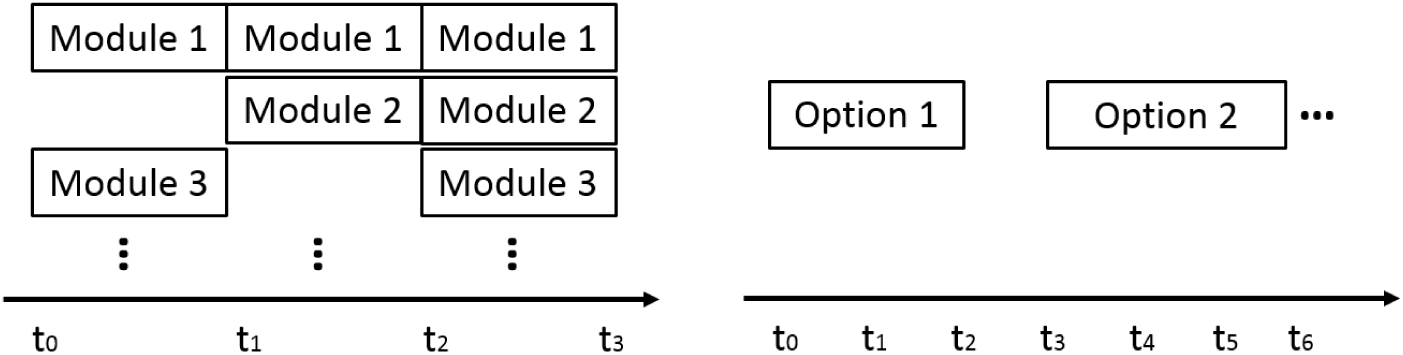
Modular reinforcement learning (left) vs. hierarchical reinforcement learning (right). Modular RL assumes modules run concurrently and do not extend over multiple time steps. Hierarchical RL assumes that a single option may extend over multiple time steps.

## 5 Discussion

This paper formalizes a modular reinforcement learning model for natural multitask behaviors. Modular RL is more suitable for modeling human behaviors in natural tasks while standard RL serves as a general model for reward-seeking behaviors. The two important variables in modular RL are module-specific reward and discount factor, which can be jointly estimated from behavioral data using the proposed modular IRL algorithm. A computer simulation demonstrated the validity and sample efficiency of the modular IRL. In a virtual-reality human navigation experiment, we showed multitask human navigation behaviors, across subjects and under different instructions, can be modeled and reproduced using modular RL.

Modular RL/IRL makes it possible to estimate the subjective value of particular human behavioral goals. Over the last 15 years it has become clear that the brain’s internal reward circuitry can provide a mechanism for the role of tasks on both gaze behavior and action choices. It is thought that the ventromedial prefrontal cortex and basal ganglia circuits encode the subjective values driving behavior [46-48]. The present work shows that it is possible to get a realistic estimate of the subjective value of goals in naturalistic behavior, and these values might reflect the underlying reward machinery. Many of the reward effects observed for neurons have very simple choice response paradigms. Thus it is important to attempt to link the primary rewards used in experimental paradigms and the secondary rewards that operate in natural behavior. Previous human experiments have typically used simple behaviors with money or points as rewards. In our experiment we used instructions to bias particular aspects of basic natural behavior with no explicit rewards.

The results provide support for a modular cognitive architecture when modeling natural visually guided behaviors. Modularization reduces the size of state space and alleviates the curse of dimensionality. Consequently modular IRL is more sample efficient than the standard Bayesian IRL. In addition, modular RL estimates a discount factor for every module hence it is more flexible and powerful than a standard RL model in which the discount factor is unitary and fixed. The modeling result suggests having such flexibility is indeed helpful. It may also explain why basal ganglia has the mechanism to implement multiple discount factors [16].

The decomposition of global task also allows humans to reuse a learned module later in a new environment. This claim is supported by the observation that task-relevant module rewards and discount factors are stable and generalize to a different task condition. When immersed in a new environment, the simple form of Eq (4) allows value function to be computed with reasonable cognitive load. It is possible that subjects learn stable values for the costs of particular actions like walking and obstacle avoidance and these subjective values factor into momentary action decisions [1]. For example, humans direct gaze to nearby pedestrians in a simple uninstructed walking context with a probability close to 0.5, with small variability between subjects [49] and a similar gaze distribution was found in a virtual environment [50]. These values may change in more complex contexts, as in the decoy effect for example [51]. The present work provides a way of testing the circumstances in which such subjective values might change.

Modular RL allows intuitive interpretation for multitask behaviors, where relative importance and reward discounting rates can be compared between modules directly. We expect this modular approach of RL can be applied to and can explain many natural tasks. [52] has shown that a wide range of human behaviors can be modeled as consisting of microbehaviors, so many behaviors are a mixture of simple modules and could potentially be modeled in this way.

A question remains of how these modules are formed originally. The intuition for a modularized strategy comes from two conjectures: learning is incremental and attentional resource is limited. From a developmental perspective, a complicated natural task is often divided in to subtasks when learning happens, e.g., curriculum learning [53], hence a real-time decision-making rule is likely to be a combination of pre-learned subroutines. A subtask is attended when needed to save computational resource.

### Limitations of the model and future work

Although modular RL/IRL is able to produce trajectories that are similar to human behavior, the match was imperfect as demonstrated by the angular difference. One difficulty with modeling human behavior is that we defined the state space and a set of modules by hand without knowing the actual state representation or task decomposition that the human uses. This may account for the discrepancy between the human and agent policies. Ideally, we could learn state representation from data, but this involves the challenging task of combining representation learning and IRL. The work in [54] provides a potential method for inferencing goals and states for the modules. Recent development in deep reinforcement learning [55] may possibly lead to a data-driven approach to IRL that can learn state representation from data.

An important assumption about the centralized arbitrator of the modules needs to be examined more carefully in the future: In our model, an agent forms global Q-values by summing up module Q-values [21,29]. There has been work examining more sophisticated mechanisms for global decision making [56,57]. For example, one could schedule modules according to an attention mechanism [56,58]. Whether these mechanisms can better explain human behaviors remains an open question that should be explored.

An important consequence of being able to get a quantitatively estimated subjective reward and discount factor of a module is that it is possible to test whether these values are stable across contexts. For example, the value of avoiding an obstacle should be stable across moderate variations in the environment such as the changes in obstacle density or changes in the visual appearance of the environment. If this is true, then it is possible to make predictions about behavior in other contexts using learned modules. And it would also be possible to use the prediction error to indicate that other factors need to be considered.

Estimates of the value of the underlying behaviors will also allow prediction of the gaze patterns subjects make in the environment. It has been suggested that gaze patterns reflect both the subjective value of a target and uncertainty about task-relevant state [2, 4, 59, 60]. For example, gaze should be frequently deployed to look at pedestrians in a crowded environment since it is important to avoid collisions and there is high uncertainty about their location. Also gaze is deployed very differently depending on the terrain and the need to locate stable footholds, reflecting the increased uncertainty of rocky terrain [61]. Estimates of the subjective value might thus allow inferences about uncertainty as well.

In conclusion, we have demonstrated that modular reinforcement learning can plausibly account for sequences of sensory-motor decisions in a natural context, and it is possible to estimate the internal reward value of behavioral components such as path following, target collection, and obstacle avoidance. The estimated reward values and discount factors enabled us to predict long walking trajectories in a novel environment. This framework provides a potentially useful tool for exploring the task structure of natural behavior, and investigating how momentary decisions are modulated by internal rewards and discount factors.

## Supporting Information Legends

**S2 Video. A sample video from collected human data.** The attached video file shows a typical experimental trial from the subject’s point of view, with motion tracking eye tracking enabled (the white cross). The task of this particular trial is to collect the targets, avoid the obstacles, and follow the path at the same time.

## Supporting Information

### Appendix 1: Simulation Results

Using a canonical 2D gridworld in reinforcement learning (RL) research, the goals are to empirically prove that modular IRL algorithm can estimate rewards and discount factors correctly, demonstrate its advantages over standard IRL, and show an example of sparse modular IRL. Part of the gridworld is shown in Fig 1. Different module objects are indicated by different colors and shapes. Behavioral data (state-action pair samples) are collected from a modular RL agent.

We first show that modular IRL is able to recover module rewards and discount factors correctly. The environment contains six modules each with ten objects. Three of them have positive rewards and the other three have negative rewards. 10 gridworlds are generated with random layouts of objects. The agent navigates each world for 6,000 steps. Non-sparse modular IRL (Eq (9)) is used to estimate *r*^(*1:6*)^ and *γ* ^(*1:6*)^ and we calculate the mean estimation and standard error. The results are shown in Table 1, it is evident that modular IRL is highly accurate in recovering the true rewards and discount factors given a large amount of data.

**Fig 1.**
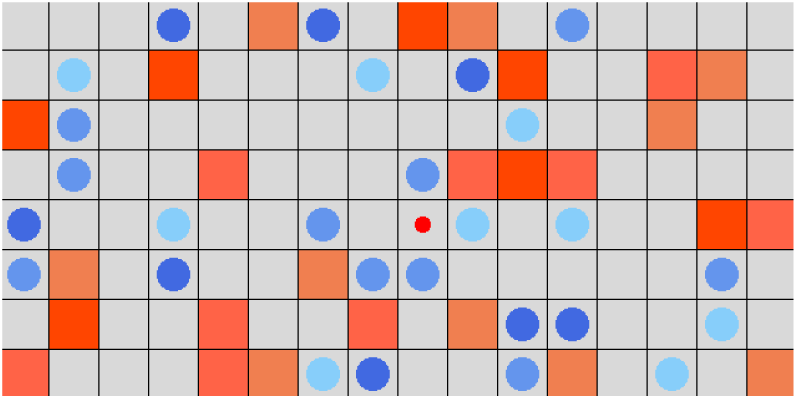
Part of the 2D gridworld test domain. Red squares are obstacles with negative reward. Blue circles are targets with positive reward. The small red dot is the modular RL agent. Different colors indicate different modules with distinct rewards and discount factors. The objects of the same module have the same color.

**Table 1.**
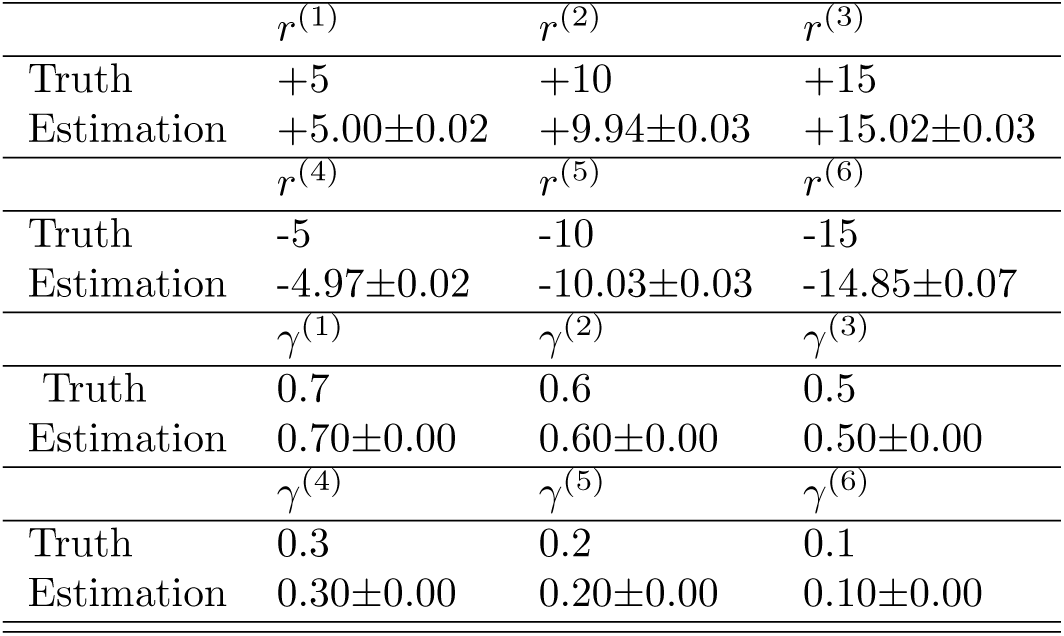
Estimated rewards and discount factors comparing to the ground truth for the six modules in the 2D gridworld experiment. The results are presented as mean ± standard error (N = 10). The estimations are highly accurate due to the availability of a large amount of data.

**Fig 2.**
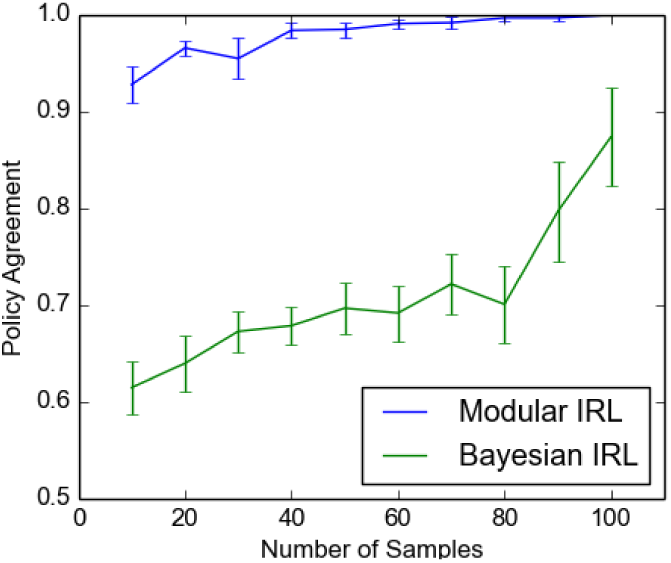
Modular IRL vs Bayesian IRL on sample efficiency, measured by average policy agreement ± standard error (*N* = 10). Modular IRL has significant higher sample efficiency.

**Modular vs. Bayesian inverse reinforcement learning** In modeling natural human behaviors, one particularly important aspect of a machine learning algorithm is its sample efficiency, given that it could be expensive to collect behavior data unlike in computer simulation. The performance of modular IRL on sample efficiency is compared with a standard non-modular Bayesian IRL [36]. We use a Laplacian prior in Bayesian IRL since the rewards are sparse. Fig 2 shows the results. The test environment has 4 modules and each has 4 objects which is made smaller because Bayesian IRL is computationally expensive. Both algorithms are given different amount of samples (state-action pairs) for training. Then policies generated using the learned rewards are compared. Policy agreement is defined as the proportion of the states that have the same policy as the ground truth, which is used because the outputs of these two algorithms are weights and rewards that can not be directly compared. Modular IRL obtained nearly 100% policy agreement with far fewer data samples compared to the Bayesian IRL.

**Sparse modular inverse reinforcement learning** Next we evaluate the performance of sparse modular IRL algorithm (Eq (10)) in terms of sample efficiency. Again the gridworld contains 10 modules and each has 10 objects. The agent only considers 2 modules, i.e., the agent makes decision by treating all other modules to have zero rewards. Therefore, the hypothetical module set has size | ℋ| = 10 and actual module set has | ℋ’| = 2.

**Fig 3.**
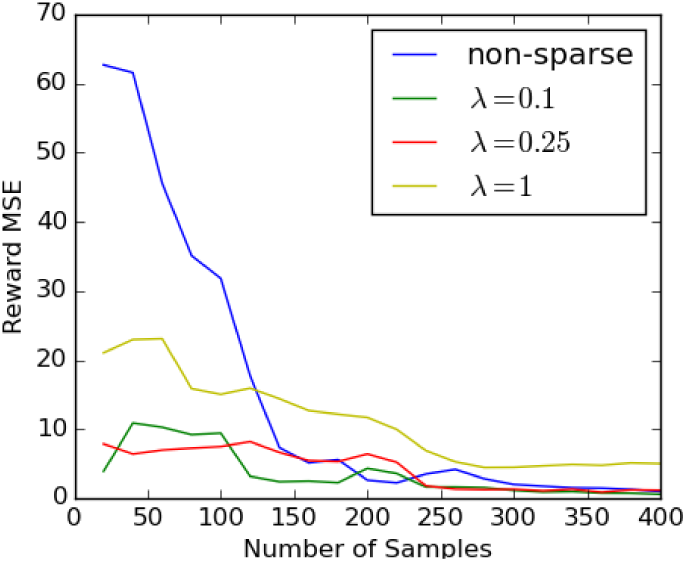
Modular IRL vs sparse modular IRL on sample efficiency, measured by mean squired error (MSE) of estimated reward. Sparsity can greatly improve sample efficiency with a carefully chosen value of λ.

**Table 2.** One-way ANOVA for individual differences in reward between subjects and across task instructions. Between-subject differences for all modules are significant in all task conditions.

| | Target $r$ | Obstacle $r$ | Path $r$ |
| --- | --- | --- | --- |
| Task 1 | $F(25, 4) = 6.53$<br>$p = 3.38 \times 10^{-11}$ | $F(25, 4) = 5.60$<br>$p = 1.16 \times 10^{-9}$ | $F(25, 4) = 4.57$<br>$p = 8.44 \times 10^{-8}$ |
| Task 2 | $F(25, 4) = 8.09$<br>$p = 1.41 \times 10^{-13}$ | $F(25, 4) = 12.11$<br>$p = 1.18 \times 10^{-18}$ | $F(25, 4) = 12.12$<br>$p = 1.16 \times 10^{-18}$ |
| Task 3 | $F(25, 4) = 7.65$<br>$p = 6.11 \times 10^{-13}$ | $F(25, 4) = 5.91$<br>$p = 3.50 \times 10^{-10}$ | $F(25, 4) = 3.17$<br>$p = 4.50 \times 10^{-5}$ |
| Task 4 | $F(25, 4) = 21.38$<br>$p = 6.57 \times 10^{-27}$ | $F(25, 4) = 5.03$<br>$p = 1.21 \times 10^{-8}$ | $F(25, 4) = 7.20$<br>$p = 3.00 \times 10^{-12}$ |

The mean squared error (MSE) of the estimated reward is shown in Fig 3. If data is scarce, the sparse version of modular IRL algorithm (λ = 0.1, 0.25) can recover rewards more accurately than the non-sparse version. Sparse modular IRL correctly identifies modules that the agent paid attention to, indicated by low MSE values obtained. As the regularization constant λ controls the importance of the regularization term, a very large λ introduces a large bias in estimation and may fail to converge to the truth, as shown by λ = 1. One can use the standard cross-validation techniques in choosing the value for λ.

### Appendix 2: One-way ANOVA for Estimated Rewards

Table 2 shows ANOVA results for individual differences in reward between subjects. Fig 4 visualizes the effect of task condition on reward function for each individual subject.

**Fig 4.**
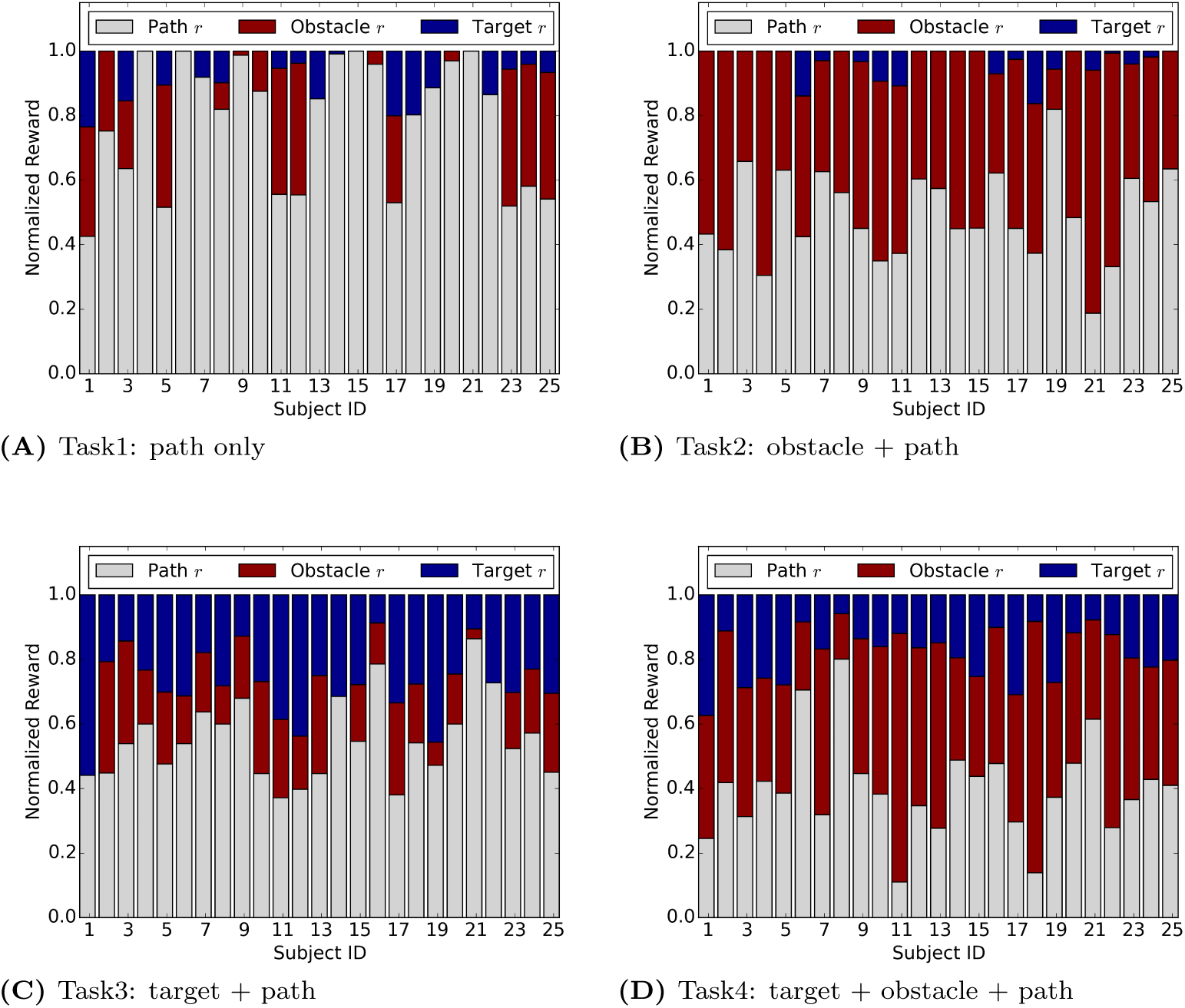
Average normalized rewards for each subject under different task instructions. The relative reward magnitude changes between tasks and agrees with task instructions. Under the same task instruction, individual differences in reward function are shown.

### Appendix 3: Bayesian Inverse Reinforcement Learning

Bayesian IRL leverages demonstrated state-action pairs, treats them individually as evidence for the underlying reward function and therefore is able to express the likelihood of reward functions given demonstrations. The normalizing factor for computing the probability of reward functions is hard to compute, hence Bayesian IRL instead adopts a Monte Carlo Markov Chain (MCMC) sampling method to acquire a set of reward samples using the unnormalized likelihood function [36]. In order to compute the likelihood of a given reward function during sampling, it is required to compute the Q-values for all the state-action pairs in the demonstration set, which means solving a reinforcement learning (RL) problem given the Markov Decision Process (MDP). Therefore, Bayesian IRL is indeed a very computationally expensive algorithm.

In order to make our human experiment environment tractable by Bayesian IRL, the virtual room is discretized into a 2D gridworld of size 32 × 24 with 0.2 × 0.2 m^2^ cells. Each cell is a state in the MDP. The actions are discretized into 8 directions so that an agent can move to any adjacent state in the gridworld. The (center) location of targets, obstacles and waypoints are treated as different feature points, which contribute to each state’s feature by distance. The problem is formulated as learning the weights for the three different features: targets, obstacles and waypoints. The three features are represented using three different continuous values at each state. More specifically, the closer a state is to an target/obstacle/waypoint, the higher the feature value for the particular object at that state. The reward at any given state is computed as the linear combination of these features using their corresponding weights. The observations are a set of state-action pairs extracted from the human’s trajectory, which are fitted to the discretization of the space.

The parameters for Bayesian IRL are set empirically. The confidence factor *α* is set at 80 and the chain length is set to be 3000 (since there are only three values, i.e. feature weights, to be tweaked, which is relatively small). A value of 0.5 is used as the discount factor for MDPs with the assumption that the decision making process of humans tends to prefer immediate rewards.

